# Preservation and Clonal Behavior of Extrachromosomal DNA in Patient-Derived Xenograft Models of Childhood Cancers

**DOI:** 10.1101/2025.08.14.670153

**Authors:** Rishaan Kenkre, Jon D. Larson, Owen S. Chapman, Jens Luebeck, Yan Yuen Lo, Megan Paul, Wenshu Zhang, Vineet Bafna, Robert J. Wechsler-Reya, Lukas Chavez

## Abstract

Extrachromosomal DNA (ecDNA) is a powerful oncogenic driver linked to poor prognosis in pediatric cancers. Whole-genome sequencing of 338 patient-derived xenograft (PDX) samples and 127 matched primary tumors across multiple childhood cancer types was used to compare ecDNA prevalence, sequence conservation, and clonal dynamics. ecDNA in PDX models frequently mirrored oncogene amplifications observed in patient tumors (e.g., MYCN, MYC, MDM2) and showed high sequence conservation. Medulloblastoma and neuroblastoma PDXs exhibited significantly higher ecDNA prevalence, consistent with strong selection or de novo formation during tumor propagation. Although ecDNA copy numbers were generally preserved, some neuroblastoma PDXs displayed marked MYCN copy gains. Single-cell multiome profiling revealed that ecDNA-positive clones either persisted or expanded dramatically in PDXs, in one case growing from a minor subpopulation to nearly all tumor cells. These findings establish PDX models as valuable systems for ecDNA research and underscore the selective growth advantage conferred by ecDNA during tumor evolution.

## INTRODUCTION

Extrachromosomal DNA (ecDNA), also known as double minutes, is a critical oncogenic driver in many different types of cancer [^1,2,3,4^]. Childhood cancer patients whose tumors contain ecDNA have significantly worse 5-year survival than patients whose tumors have chromosomal or no amplifications [^3,4,5^]. Due to the absence of centromeres, ecDNA exhibits random inheritance, driving copy number amplification, promoting intratumoral heterogeneity, accelerating tumor evolution, and contributing to treatment resistance [^6^]. These features emphasize the importance of studying the role of ecDNA during tumor development, recurrence, and evolution.

Patient-derived models, such as cell lines and xenograft (PDX) mouse models, are widely used to study tumor biology and therapeutic vulnerabilities. Pharmacological inhibition studies in PDX mouse models are important preclinical experiments for evaluating tumor responses *in vivo*. PDX models largely recapitulate the histological features, DNA methylation profiles, and intratumoral heterogeneity of the tumors from which they were derived [^7,8^]. However, the behavior of ecDNA during PDX model development and propagation has not yet been analyzed. To evaluate the representation of ecDNA in PDX models compared to the tumors from which the models were derived, this study analyzes the occurrence and preservation of ecDNA sequences and copy numbers in PDX models compared to human tumors in a broad spectrum of childhood solid cancers.

## RESULTS

### ecDNA in human tumors and PDX models amplify the same oncogenes

To investigate ecDNA in PDX models of childhood cancers, we accessed whole-genome sequencing (WGS) data from St. Jude Cloud’s Childhood Solid Tumor Network [^9,10^], a medulloblastoma (MBL) cohort from Rady’s Children’s Hospital, San Diego, and other sources [^3^] **(Figure 1)**. Collectively, this dataset included 338 PDX samples derived from 288 patients across 31 childhood cancer types. In some cases, multiple PDX models were descended from the same patient tumor, leading to more than one PDX sample per patient in the dataset (**Supplementary Table 1**). By employing AmpliconArchitect (AA) [^10^] and AmpliconClassifier (AC) [^11^], we identified 175 ecDNA sequences in 106 PDX models (31%) derived from 93 patients across 14 cancer types (**Figure 2a, Supplementary Table 2**).

**Figure 1:**
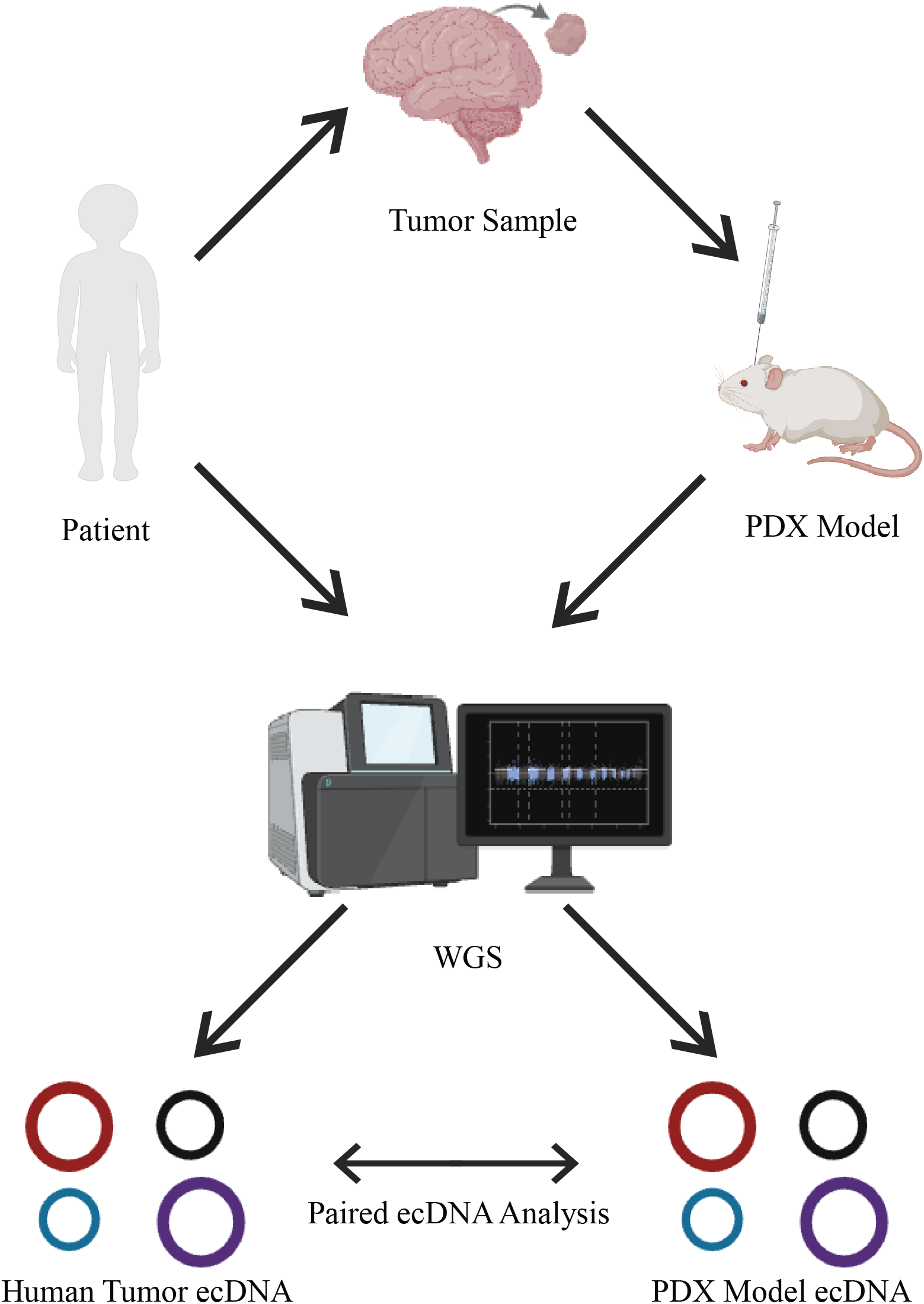
Study overview. Graphical abstract of the study design displaying each stage from tumor sample extraction to paired ecDNA analysis.

**Figure 2:**
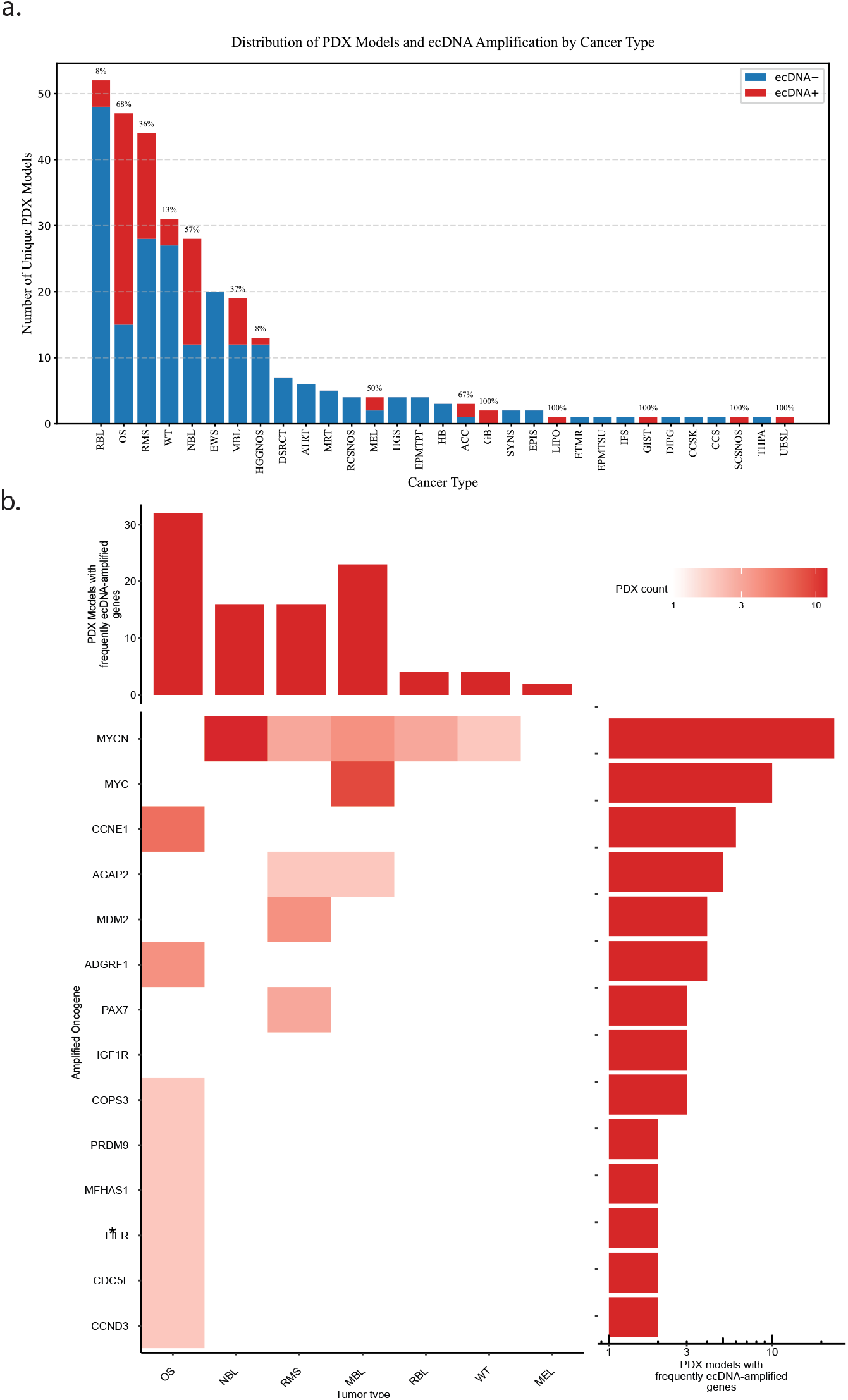
Distribution of PDX models across childhood cancer types. **(a)** Blue bars show the number of PDX models by cancer type. Red bars show the number of ecDNA-positive PDX models by cancer type. (**b)** Heatmap indicates the number of PDX samples with ecDNA amplification of a given gene in that tumor type. The star indicates oncogene found in the PDX model cohort but not in the pedpancancer human Tumor cohort. The top barplot indicates the number of PDX models with ecDNA sequences amplifying any of the indicated genes for each cancer type. The right barplot indicates the number of ecDNA sequences amplifying the indicated gene across all tumor types.

In this cohort of 106 ecDNA-positive PDX models, a total of 14 different oncogenes wee recurrently amplified on ecDNA (**Figure 2b**). As in primary childhood cancers [^5^], *MYCN* was the most frequently amplified oncogene with occurrence in 24 PDX models derived from neuroblastoma (NBL), rhabdomyosarcoma (RMS), Wilms tumor (WT), retinoblastoma (RBL), and MBL[^11,12,13^]. Furthermore, *MYC, CCNE1, MDM2*, and *IGF1R*, showed consistent amplification patterns in both PDX models and human tumors, with *MYC* most prevalent in MBL, *CCNE1* and *IGF1R* most prevalent in OST, and *MDM2* most prevalent in RMS (**Figure 2b**). Notably, *LIFR* amplification was detected exclusively in the PDX cohort but not in the reference human tumor population of childhood cancer patients (N = 3,631) [^5^]. *LIFR* serves a role as both a tumor promoter and suppressor, influencing tumor growth and development in various cancers, partly through the regulation of p53 expression [^14,15^]. When compared to this reference cohort of childhood cancer patients [^5^], we observe a statistically significant increase of ecDNA in this cohort of PDX tumors (9% vs. 31% ecDNA-positive human tumors or PDX models, respectively, **Fisher’s exact test, p < 0.001)**. Overall, these analyses demonstrate that ecDNA in PDX models of childhood cancers largely recapitulates the oncogene amplifications observed in human tumors. The persistent amplification of key oncogenes such as *MYCN* and *MDM2* across both PDX models and primary tumors validates the biological relevance of these models in investigating ecDNA-driven oncogenesis.

### ecDNA amplifications are enriched in PDX models

To evaluate the frequency of ecDNA in PDX models compared to the reference cohort of primary human tumors, we analyzed seven pediatric cancer types for which at least five PDX models were available (262 PDX models in total). Statistical analysis revealed enrichment of ecDNA detected in PDX compared to the reference human tumors in MBL and NBL (□**2 = 27.7 and 10.1, p = 1.4e-7 and 0.0015 respectively, Figure 3**). A similar but nonsignificant trend was observed in OST tumors (51%) **(Chi-squared test of independence, p = 0.110, Figure 3)**. For the remaining tumor types, no difference in the frequency of ecDNA was observed comparing PDX models and human tumors.

**Figure 3:**
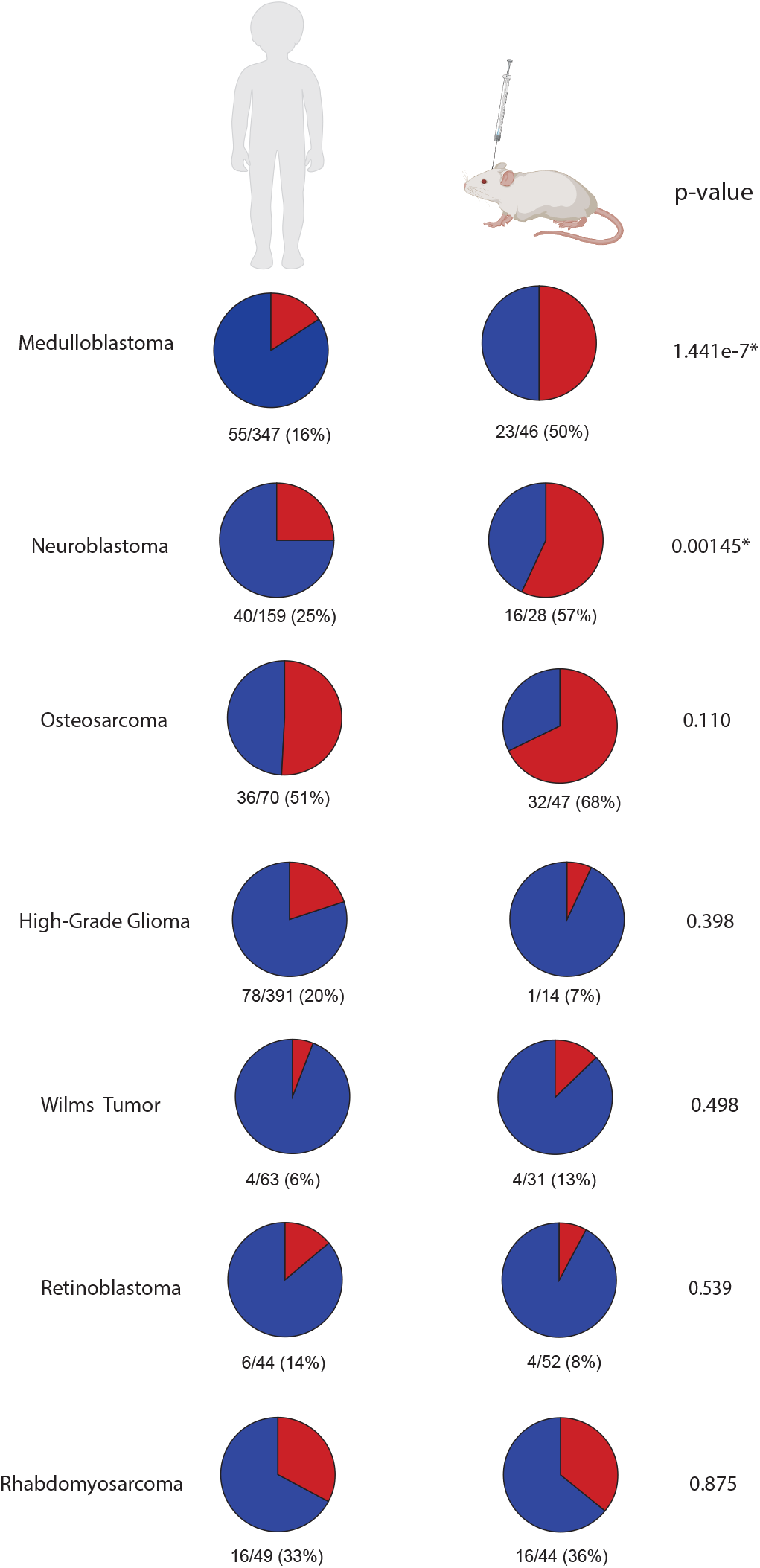
Frequency of ecDNA in human tumors and PDX models. Pie charts displaying ecDNA prevalence in primary human tumors of different childhood cancer types as analyzed in a previous pediatric pan cancer study of ecDNA (left column) compared to PDX models analyzed in this study (right column). Red = ecDNA-positive, blue = ecDNA-negative. Statistical analysis shows significant enrichment of ecDNA prevalence in PDX tumors for medulloblastoma and neuroblastoma (Chi-squared test of independence p-values shown for each tumor type in the right columns). Similarly, an increased frequency of ecDNA was observed in PDX models derived from osteosarcoma patients; however, enrichment was below significance.

Enrichment of ecDNA in PDX models compared to the patient reference cohort is challenging to interpret due to potential differences in sampling methodology and tumor selection criteria between the two cohorts. To enable a direct comparison, we next focused our analysis on 127 patients for whom WGS data are available for both the primary human tumors and the PDX models derived from these tumors **(Supplementary Table 3)**. In the majority of cases (106/127, 83%), ecDNA status remained unchanged in the PDX models compared to the primary tumors (**Figure 4**). This included 30 out of 33 patients (91%) whose tumors remained ecDNA-positive, with only 3/33 (9%) cases losing ecDNA in PDX models. Notably, 19/94 (20%) ecDNA-negative primary tumors gained ecDNA in their corresponding PDX models **(Figure 4, Supplementary Table 3)**. Consequently, ecDNA was more prevalent in the PDX models than in their corresponding human tumors **(McNemar’s test, *p* = 0.0014)**. Among the 94 ecDNA-negative primary tumors, two distinct patterns emerged based on their amplification status. For the 16 tumors with chromosomal amplifications, half (50%) of their PDX models converted to ecDNA amplifications, while 38% retained chromosomal amplifications and 13% lost amplifications entirely. For the 78 primary tumors without any amplifications, most PDX models (65%) remained unamplified, although some gained chromosomal (21%) or ecDNA (14%) amplifications.

**Figure 4:**
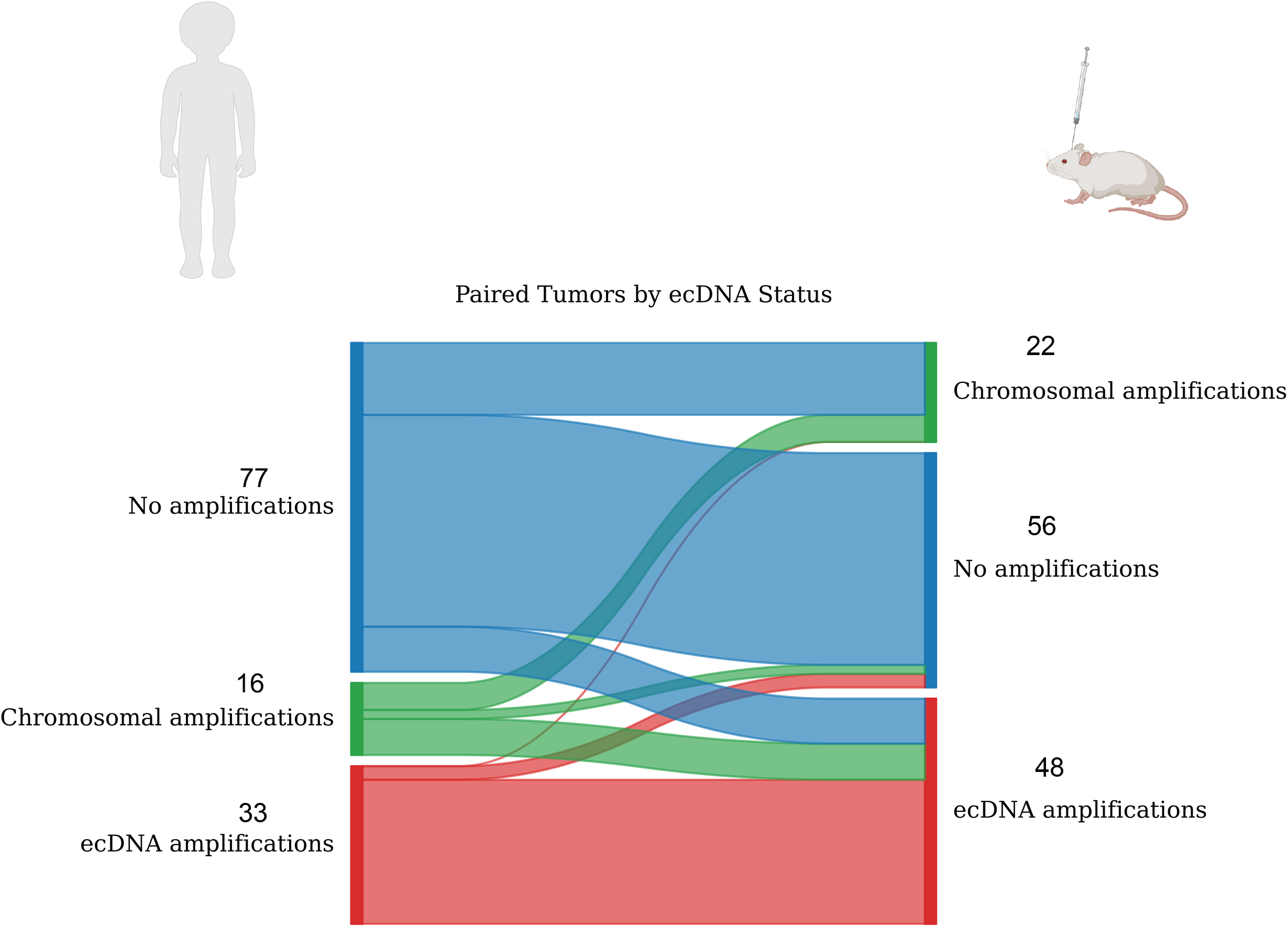
Comparison of DNA amplification status in human tumours and their PDX models. Sankey diagram that displays the flow of ecDNA status from human tumors to the corresponding PDX models for 127 pairs. Primary tumors on the left side and PDX models on the right side. Blue flows indicate primary tumor is ecDNA-negative with no other amplifications; green flows indicate primary tumor is ecDNA-negative with chromosomal amplifications; red flows indicate primary sample is ecDNA-positive.

To analyze potential technical causes of differences in ecDNA status between PDX models and human tumors, we visually inspected WGS coverage profiles at the genomic ecDNA loci. First, we examined the three PDX models for which no ecDNA was detected in the PDX tumors, even though the human tumors from which they were derived were ecDNA-positive. In two of the three PDX models, increased WGS coverage was observed at the ecDNA loci, suggesting that ecDNA amplifications may be present in the PDX models but remained below the detectable sequencing coverage threshold (**Supplementary Figure 1)**. We next evaluated the 18 cases that exhibited *de novo* ecDNA emergence in the PDX models that were not observed in the corresponding human tumor. Interestingly, 13 of the 19 (68%) PDX models with newly acquired ecDNA were either NBL (n=4) or OST (n=9), despite these tumor types comprising a smaller proportion (38/127, 30%) of the cohort with paired samples analyzed in this study **(Supplementary Table 4)**. One of the three NBL PDX models that acquired ecDNA-*MYCN* amplifications had a chromosomal MYCN amplification in the human tumor **(Supplementary Figure 2a)**, whereas the other two did not have visible traces of WGS enrichment in the human tumors at the *MYCN* locus **(Supplementary Figures 2b–c)**. Six of the nine OST PDX models that gained ecDNA showed chromosomal amplifications in the human tumors, whereas the other three models did not show any traces of WGS enrichment (see **Supplementary Figure 2d** for case SJOS012409). While more research is required, these observations suggest that the larger number of ecDNA-positive PDX tumors compared to human tumors as observed for some tumor types (**Figure 4**) might be caused by *de novo* emergence, or by the outgrowth of an undetectable ecDNA+ clone, during the development of PDX models.

Most tumor types can be stratified into molecular subgroups with distinct clinical and molecular characteristics. For example, in a previous study of 468 MBL patients [^3^], we observed a varying frequency of ecDNA in the different MBL subgroups where ecDNA was detected in 27% of SHH, 18% of Group 3, and 14% of Group 4 MBL tumors, while being absent in WNT tumors **(Figure 5a)** [^3^]. While the available cohort of PDX models is not large enough for a representative stratification of all childhood tumor types into their molecular subgroups, there is a sufficiently large number of 46 MBL PDX models representing the four molecular MBL subgroups. As in human tumors, no ecDNA was observed in MBL WNT PDX models, while a substantial amount of SHH (67%), Group 3 (56%), and Group 4 (25%) MBL PDX models contained ecDNA **(N = 46, Figure 5b)**. Thus, the enrichment of ecDNA in MBL PDX models compared to human MBL patients was concentrated on PDX models of the SHH MBL and Group 3 MBL subgroups **(Fisher’s exact test, p < 0.002 for both subgroups)**. When comparing genes amplified on ecDNA in human MBL tumors and MBL PDX models, we observe 13 oncogenes consistently amplified across both PDX models and patient tumors, including *MYC* and *MYCN* that are commonly amplified on ecDNA in MBL and other tumor types (**Figures 5c, d**).

**Figure 5:**
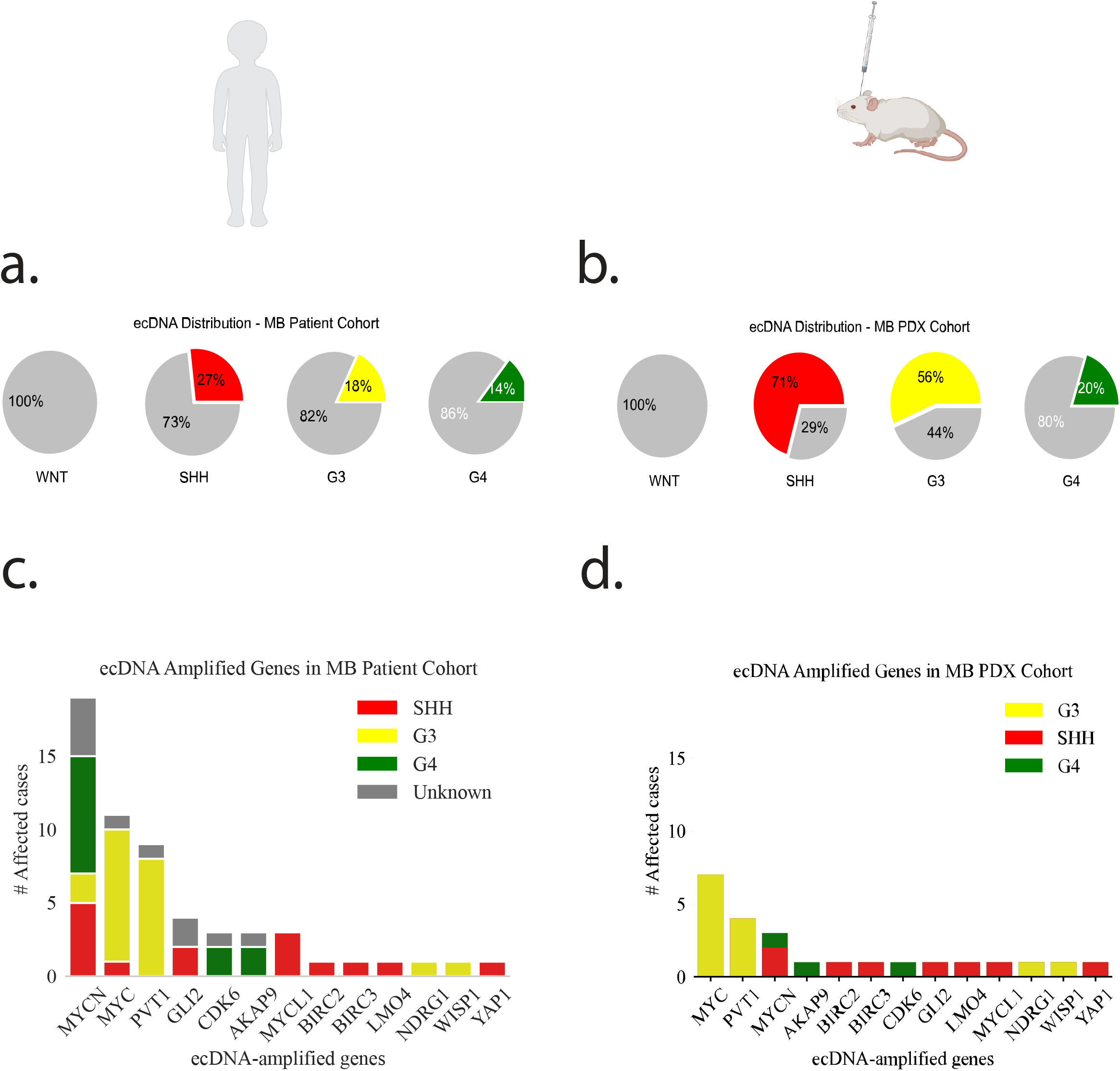
Frequency of ecDNA in human tumors and PDX models across medulloblastoma subgroups. **(a)** Frequency of ecDNA-amplified genes in the MB PDX model cohort grouped by molecular subgroup **(b)** Frequency of ecDNA-amplified genes in the MB PDX model cohort grouped by molecular subgroup **(c)** Frequency of ecDNA in the MB patient cohort grouped by molecular subgroup. In the MB human tumor cohort, the distribution of ecDNA-positive cases across molecular subgroups was as follows: WNT (0/24), SHH (30/112), G3 (19/107), and G4 (26/181). **(d)** Frequency of ecDNA in the MB patient cohort grouped by molecular subgroup. In the MB PDX cohort, the distribution was: WNT (0/3), SHH (12/17), G3 (9/16), and G4 (2/10).

### ecDNA sequence is largely conserved in PDX models

To assess whether ecDNA sequences are conserved in PDX models compared to their tumors of origin, we employed a metric derived from Jaccard similarity [^16,17^] to calculate DNA sequence similarities of ecDNA amplifications **(see Methods)**. We identified 45 ecDNA sequences from 30 tumor-PDX pairs in which both the human tumors and the PDX models were ecDNA-positive, allowing a direct comparison of ecDNA sequence similarities. **(Supplementary Table 5)**. Among the 30 ecDNA-positive human tumor-PDX pairs, four (13%) showed no overlap in amplicon coverage, indicating that ecDNA amplifications arose from entirely different genomic loci in the tumor and PDX models. The lack of overlap suggests that, in these cases, the PDX models may have developed from tumor clones harboring distinct ecDNAs, rather than directly mirroring the ecDNA architecture of the original tumor. In the remaining 26 human tumor-PDX pairs, we observed 4/45 (8.9%) ecDNAs in the human tumors that only partially overlapped with ecDNAs in the PDX models (amplicon overlap < 95%) and 38/45 (84.4%) ecDNAs originating from near-identical genomic loci (amplicon overlap > 0.99; **Figure 6a**).

**Figure 6:**
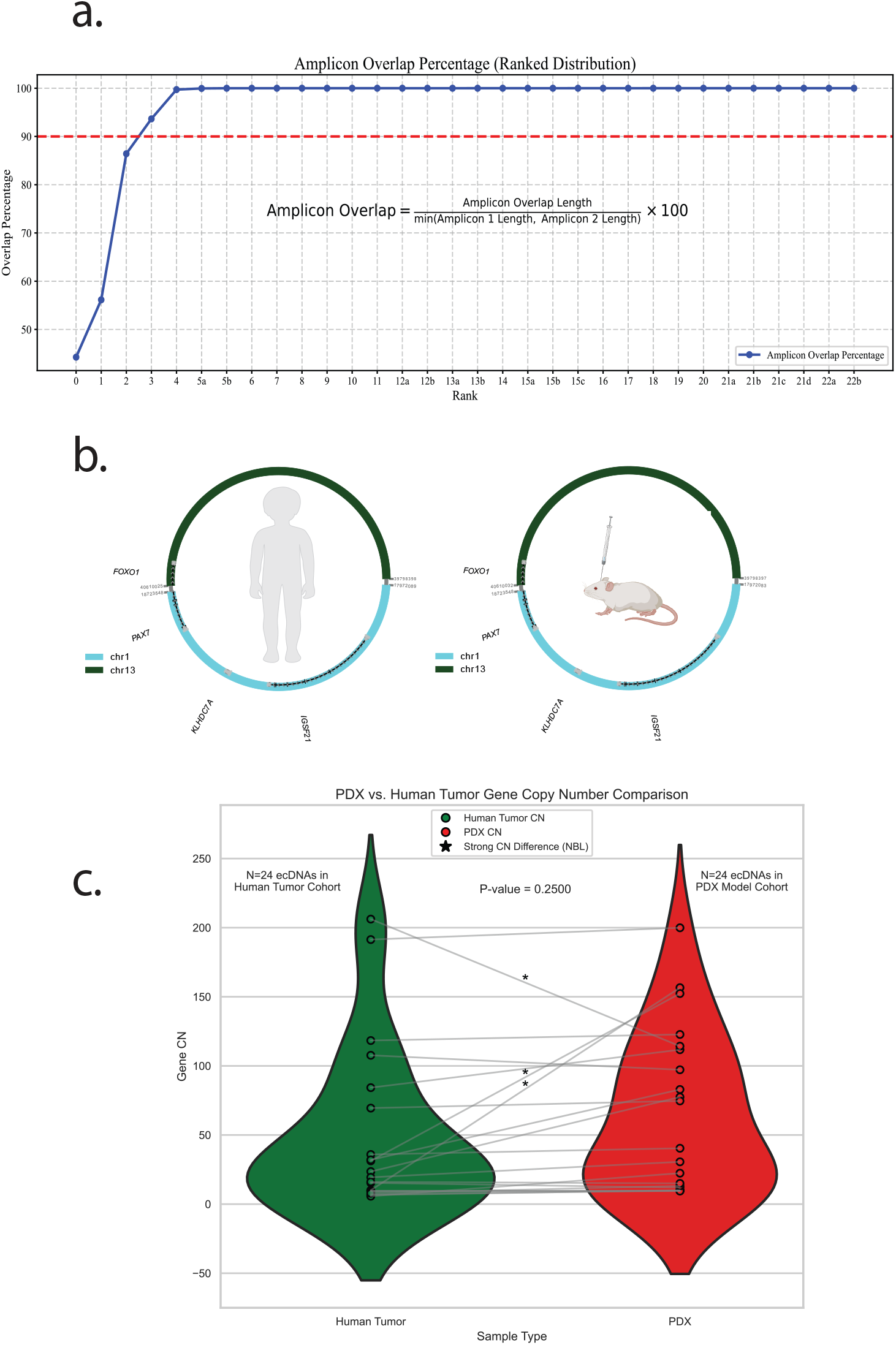
Amplicon and copy number similarity of ecDNA-positive human tumors and their PDX models. **(a)** Ranked line chart of amplicon length coverage percentage between ecDNA-positive amplicon amplicons in human tumor-PDX model pairs. Red line cutoff for pairs with amplicon overlap coverage at 90%. **(b)** Circos circular visualizations of patient SJRHB010468 named SJRHB010468_D1 and SJRHB010468_X1, respectively. **(c)** Paired copy number comparison violin plot after removing pairs with low amplicon length coverage percentage. Contains 23 human tumors and 23 PDX models and 24 paired ecDNAs. Removed samples of patients SJNBL013763, SJRHB011, SJOS001126, SJOS013768, SJOS030605, SJRHB063823, and Med1911FH.

We next evaluated amplicon similarity scores that quantify shared features between focal amplifications based on genomic composition and breakpoint locations. Applying this approach to the 26 tumor-PDX pairs with overlapping ecDNAs, we observed distinct levels of conservation **(see Methods)**. Very high amplicon similarity scores exceeding 0.995 were observed in 6/26 (23%) tumor–PDX pairs (**Supplementary Figure 3a**). Two tumor-PDX pairs exhibited complete sequence conservation (similarity score = 1.0): an ARMS case (SJRHB010468) with conserved amplification of *FOXO1, IGSF21, KLHDC7A*, and *PAX7* on Chromosome 13 **(Figure 6b)**, and an OST case (SJOS016016) with preserved amplicons on Chromosome 1, including *MCL1, ARNT, MLLT11*, and *SETDB1* **(Supplementary Figure 4a)**. However, the range of similarity scores was between 0.11-1 with a mean of 0.66 **(Supplementary Figure 3a)**. These lower similarity scores are largely driven by lower segment and breakpoint scores averaging at 0.87 and 0.58, respectively (**Supplementary Figure 3b, c)**. Examples of moderate to high conservation included an OST pair (SJOS001121) with a similarity score of 0.84 **(Supplementary Figure 4b)**, and an NBL case (SJNBL124) with a score of 0.74, despite breakpoint variability **(Supplementary Figure 4c)**. A divergent NBL case (SJNBL046148) showed minimal structural similarity (score 0.37), largely due to low breakpoint concordance (0.034) **(Supplementary Figure 4d)**. The low breakpoint concordance highlights breakpoint divergence as a key contributor to structural variability. These findings indicate greater conservation of amplified sequences than breakpoint structures, suggesting that recombination of ecDNA-amplified sequences occurs during or after PDX model establishment. Overall, these findings demonstrate that the majority of ecDNA-amplified sequences are conserved at the sequence level, with structural variability arising predominantly from breakpoint divergence rather than wholesale loss of amplified content.

### PDX models recapitulate the ecDNA copy number observed in human tumors

We next investigated whether the copy number of (onco-) genes amplified on ecDNA is conserved in PDX models compared to the human tumors from which they are derived. Here, we examined 24 ecDNAs from 23 primary tumors that have ecDNA-amplified genes in both human primary tumors and PDX models and whose ecDNAs have an amplicon overlap coverage of ≥ 90%. While the average ecDNA copy number was higher in the PDX models than in their matched primary tumors, this difference was not statistically significant (Wilcoxon signed-rank test, p = 0.25; **Figure 6c**). Overall, ecDNA copy numbers appeared to be relatively stable between human tumors and PDX models, with a few notable outliers. For example, two NBL tumors (SJNBL013762 and SJNBL030824) showed pronounced increases in ecDNA copy number in the PDX and both harbored *MYCN*. Conversely, one *MYCN*-amplified tumor (SJNBL046148) showed a decrease in ecDNA copy number in the PDX **(Figure 6c)**. Overall, these findings demonstrate broad ecDNA copy number conservation in PDX tumors across pediatric cancers.

### ecDNA-positive cells exhibit distinct clonal dynamics during PDX development

We have previously employed multiome (ATAC/RNA) single nucleus sequencing to identify and characterize tumor cells harboring ecDNA in a primary SHH MBL tumor (‘RCMB56-ht’) [^3^]. To analyze the clonal behavior of ecDNA-positive tumor cells during PDX development, we now profiled a PDX tumor derived from RCMB56-ht (i.e. RCMB56-pdx) using the same multiome single nucleus sequencing approach. In addition, we used multiome single cell sequencing to analyze a primary G3 MBL tumor amplifying MYC on ecDNA (i.e. RC123-ht) and a PDX tumor that we derived from that patient’s tumor (i.e. RC124-pdx).

In the primary SHH MB tumor sample (RCMB56-ht), we estimated that only 8% of tumor cells contained ecDNA amplifications [^3^] **(Figure 7a, see Methods)**. Analysis of the corresponding PDX model (RCMB56-pdx) revealed that almost all cells (99.6%, 9,608 out of 9,620) contained the ecDNA amplifications present in the ecDNA-positive tumor cells of the human tumor **(Figure 7b)**. Integration and batch correction of single-cell transcriptomes (see **Methods**[^17^]) revealed substantial transcriptional similarities between RCMB56-pdx cells and the ecDNA-positive tumor cells in RCMB56-ht **(Supplementary Figure 5)**. The combined similarity of gene expression and copy number profiles suggests that the PDX tumor originated from clonal expansion of the ecDNA-positive cells in the primary tumor.

**Figure 7:**
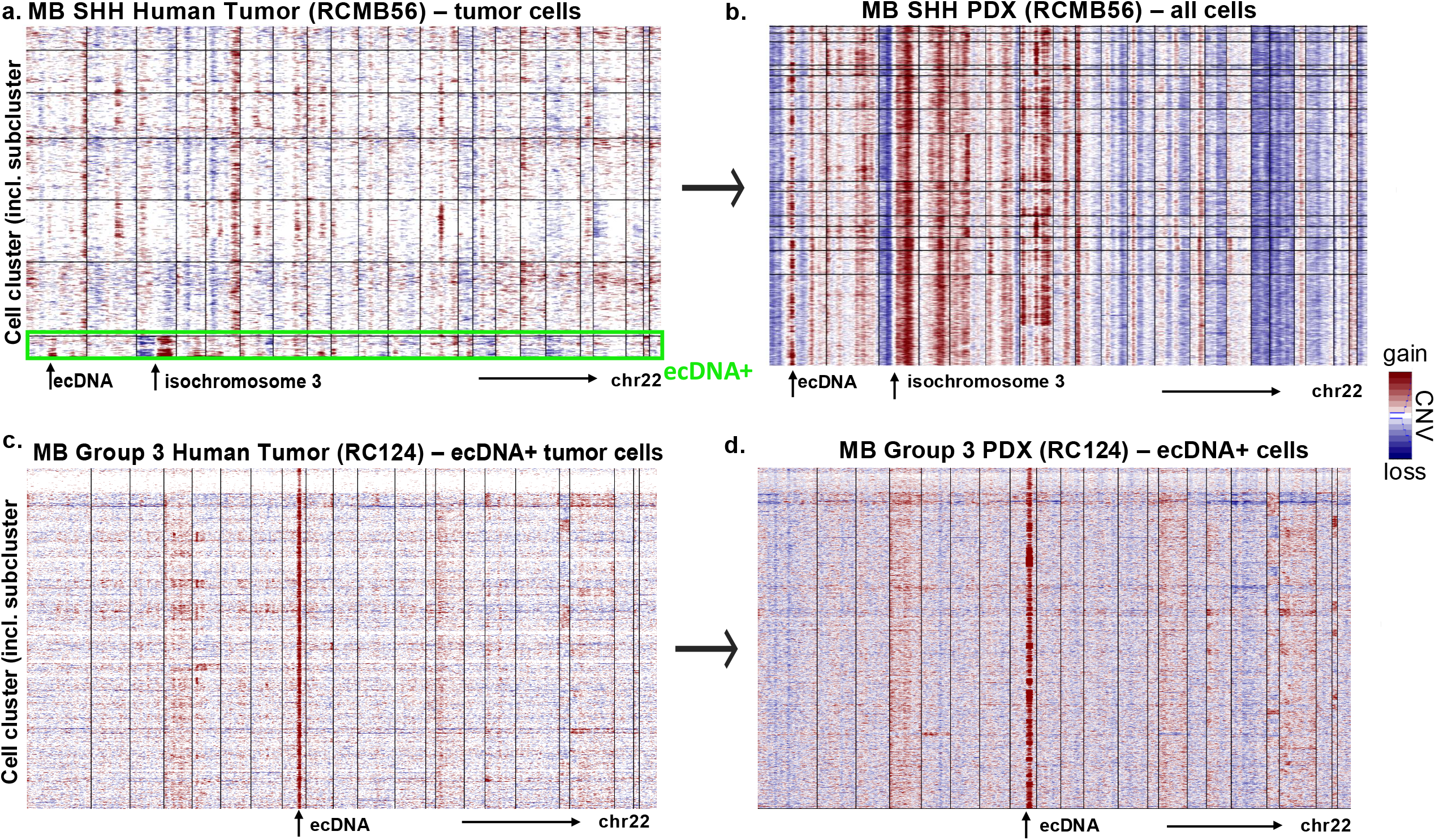
Tracking clonal dynamics of ecDNA-positive tumor cells in PDX models. **(a)** Copy-number profiles of individual cells in the MBL SHH tumor RCMB56-ht, including the 8% cDNA+ tumor cells clustered at the bottom. **(b)** Copy-number profiles of the RCMB56-pdx cells, showing that almost all cells in the PDX tumor resemble the CNV profile of the ecDNA+ tumor cells in the human tumor. **(c)** Copy-number profiles of the ecDNA-positive tumor cells of the Group 3 MBL tumor RC124-ht, representing >99% of all cells in the human tumor. **(d)** Copy-number profiles of the ecDNA-positive RC124-pdx cells, showing that almost all cells in the PDX tumor resemble the CNV profile of the ecDNA+ tumor cells in corresponding human tumor, including the MYC amplification on chromosome 8.

In contrast to the SHH MBL tumor (RCMB56-ht), ecDNA harboring the MYC oncogene was present in almost all the individual cells in the primary human G3 MBL tumor (RC124-ht) **(Figure 7c)**. Single cell analysis showed that the same ecDNA was omnipresent in the corresponding PDX tumor (RC124-pdx, **Figure 7d**), demonstrating maintenance of ecDNA throughout PDX development.

Overall, these findings revealed that ecDNA-positive tumor cells can exist either as a minor subpopulation or as a widespread cell population within primary human tumors. In RCMB56, ecDNA-bearing cells were initially rare but underwent marked clonal expansion during PDX development, suggesting a strong selective advantage for ecDNA-driven clones. Conversely, in RC124, ecDNA-positive cells were already abundant in the primary tumor and persisted as the dominant clone in the PDX. Despite these differences in initial prevalence, the resulting PDX tumors in both cases were composed almost entirely of ecDNA-positive cells. This convergence highlights the potent growth advantage conferred by ecDNA and suggests that ecDNA-bearing clones are preferentially selected during PDX establishment, regardless of their frequency in the original tumor.

## DISCUSSION

This study provides a comprehensive overview of ecDNA in pediatric cancer PDX models, highlighting its prevalence, oncogenic content, and conservation relative to patient tumors. We observed that ecDNA frequently recapitulates oncogene amplifications found in human cancers, is generally preserved during PDX establishment, and reflects subtype-specific patterns across tumor types. These findings support the utility of PDX models in studying ecDNA biology and their implications for pediatric cancer progression and treatment.

The observation that ecDNA is enriched in PDX models compared to primary tumors suggests a selective advantage for ecDNA-positive cells during xenograft establishment. This was particularly evident in neuroblastoma and medulloblastoma, where significant enrichment of ecDNA was observed in the PDX models of these tumor types. The preferential outgrowth of ecDNA-positive cells, as demonstrated in our SHH-MB RCMB56 case study in which an ecDNA-positive subclone (8% of cells) expanded to dominate the PDX model (99.6% of cells), highlights the aggressive and proliferative nature of ecDNA-harboring cells. This expansion likely reflects a selective pressure during PDX establishment, potentially favoring cells with enhanced proliferative capacity conferred by ecDNA-amplified oncogenes.

Conservation of ecDNA sequences and amplification of oncogenes between primary tumors and PDX models is encouraging for translational cancer research. With approximately 84% of ecDNA amplicons displaying substantial sequence conservation (> 90% overlap coverage), PDX models largely recapitulated the ecDNA sequences amplified in primary tumors. However, the variability of the breakpoint positions and the differences in the overall similarity scores of the amplicons indicate recombination of ecDNA during model development, although reconstruction error of ecDNA from WGS data may also play a role.

While there was a trend towards increased copy numbers of ecDNA-amplified oncogenes in PDX models compared to their human tumors of origin, this difference did not reach statistical significance, indicating that ecDNA copy numbers are generally conserved during PDX model development. The particularly dramatic copy number increases in MYCN-amplified neuroblastoma cases underscores the potential importance of this oncogene during PDX establishment.

The emergence of ecDNA-positive PDX tumors derived from some ecDNA-negative human tumors has raised important considerations for model interpretation. This phenomenon was particularly prevalent in neuroblastomas and osteosarcomas, suggesting that tumor-specific mechanisms influence ecDNA formation selection or stability. Whether these newly detected ecDNAs represent outgrowth of pre-existing minor subclones below detection thresholds in primary samples, or truly *de novo* formation during PDX establishment, remains an important question for future investigation.

Future studies should investigate the mechanisms driving the selective advantage of ecDNA-positive cells during PDX establishment and explore whether similar enrichment occurs in other model systems, such as organoids or cell lines. Understanding the factors that promote ecDNA formation and maintenance in different environmental contexts could reveal new therapeutic vulnerabilities.

In conclusion, our findings validate PDX models as valuable tools for studying ecDNA biology in childhood cancers. Longitudinal sampling during PDX tumor growth and under therapeutic pressure could provide valuable insights into the dynamics of molecular evolution, clonal selection, and ecDNA-driven therapy resistance.

## METHODS

### Data Reference

The WGS data of PDX samples and patient tumors was accessed using the St Jude Cloud Data Portal (https://platform.stjude.cloud/data/). WGS data was preprocessed and aligned according to the internal pipelines at St. Jude (hg38). Docker containers of Amplicon Architect software were installed on the DNANexus cloud genomics platform (see Methods: “ecDNA detection and classification”). In addition, low-coverage WGS data for MB PDX models and human tumors were obtained from a previous publication[^18^]. Eligibility for inclusion was determined by the availability of samples in the data portal. No attrition was recorded during the study period.

### ecDNA detection and classification from bulk WGS

To detect ecDNA, all samples in the WGS cohort were analyzed using the AmpliconSuite-pipeline[^16^] v1.1.0, AmpliconArchitect[^19^]v1.3, and AmpliconClassifier[^20^]v0.4.4, either with the GRCh38 or GRCh37 reference genomes. Briefly, the Amplicon Architect algorithm was performed as follows. Copy number segmentation and estimation were performed using the CNVkit v0.9.664. Segments with copy number ≥ 4.5 were extracted using AmpliconSuite-Pipeline (November 2023 update) as “seed” regions. For each seed, AmpliconArchitect searches for the region and nearby loci for discordant read pairs that are indicative of genomic structural rearrangements. Genomic segments are defined based on boundaries formed by genomic breakpoint locations (identified by discordant reads) and modulations in genomic copy number. A breakpoint graph of the amplicon region was constructed using CN-aware segments, and genomic breakpoints and cyclic paths were extracted from the graph. Amplicons were classified as ecDNA, breakage-fusion-bridge, complex, linear, or no focal amplification using the heuristic-based companion script AmpliconClassifier. Biological samples with one or more classifications of “ecDNA” were considered ecDNA-positive, and all others were considered ecDNA-negative.

Code is available at:

AmpliconSuite pipeline: https://github.com/AmpliconSuite/AmpliconSuite-pipeline

PrepareAA: https://github.com/jluebeck/PrepareAA

AmpliconArchitect: https://github.com/virajbdeshpande/AmpliconArchitect

AmpliconClassifier: https://github.com/jluebeck/AmpliconClassifier

### Patient metadata, survival, and medulloblastoma subgroup annotation

Where available, patient samples and models were assigned metadata annotations including age, sex, survival, and MB subgroup, based on previously published annotations of the same tumor or model[^18,21–26^]. Sample metadata are also available in some cases from the respective cloud genomics data platforms: https://dcc.icgc.org/ (ICGC), https://pedcbioportal.kidsfirstdrc.org/, https://portal.kidsfirstdrc.org/ (CBTN), and https://pecan.stjude.cloud/ (St Jude). Where primary sources disagreed on a metadata value, that value was reassigned to the NA. Patient tumors from the CBTN were assigned molecular subgroups based on the consensus of two molecular classifiers using RSEM-normalized FPKM data: MM2S69 and D3b medulloblastoma classifier at the Children’s Hospital of Philadelphia (https://github.com/d3b-center/medullo-classifier-package). To determine the molecular subgroup of PDX samples, we generated or obtained DNA methylation profiles (Illumina 450k or EPIC) from a previous publication [^22^]and classified samples by molecular subgroup according to the DKFZ brain tumor methylation classifier (https://www.molecularneuropathology.org/mnp)[^21^].

### Animals

NOD-SCID IL2Rg null (NSG) mice used for intracranial human tumor transplantation were purchased from the Jackson Laboratory (#005557). All experiments were performed in accordance with national guidelines and regulations and with the approval of the Animal Care and Use Committees at the Sanford Burnham Prebys Medical Discovery Institute and University of California, San Diego (San Diego, CA, USA).

### Establishment and maintenance of RCMB56-pdx and RC124-pdx

RCMB56-pdx and RC124-pdx were established by implanting 0.5-1×106 dissociated patient tumor cells directly into the cerebellum of NSG mice. Subsequent tumors were harvested from the mice, dissociated, and re-implanted into new NSG mice without in vitro passage. Ex vivo experiments were performed using RCMB56-pdx and RC124-pdx cells at passage 1 (p1) or higher.

### Multiome Single-Cell Seq

We generated new multiome single-cell sequencing data from the corresponding PDX model RCMB56-pdx as previously described [^3^]. Briefly, fresh PDX tumor cells were dissociated for single-cell multiome ATAC and gene expression sequencing (10X) according to the manufacturer’s instructions. Sequencing was performed using an Illumina NovaSeq S4 200 instrument at a depth of at least 250M reads for scATAC-seq and 200M reads for scRNA-seq. In addition, multiome single-cell sequencing data of the human tumor RCMB56-ht was made available by Chapman et al [^3^].

### Pairwise similarity of ecDNAs sequences

Overlapping focal amplifications were compared to quantify amplicon similarity by quantifying the relative degrees of shared overlap in genomic coordinates and SV breakpoint locations. These calculations were implemented in the feature_similarity.py script, available in the AmpliconClassifier repository (https://github.com/AmpliconSuite/AmpliconClassifier).

We defined two measurements of similarity based on Jaccard indices. JaccardGenomicSegment similarity is a Jaccard index computed using two sets formed by the coordinate ranges of genomic intervals comprising two focal amplifications. The second is Jaccard–breakpoint similarity, which is a Jaccard index computed on two sets formed by the locations of SV breakpoint junctions in the two focal amplifications. Two SV breakpoint junctions were determined to be the same if the total absolute difference between the measured genomic endpoints of the junction was less than 250bp.

The amplicon feature similarity script supports the comparison of amplicons globally for all amplicon regions, but can also be run in a restricted mode, which limits the comparison to specific regions of the genome. Furthermore, as AmpliconArchitect may include flanking regions that are not focally amplified as part of the amplification itself, the amplicon similarity script filters from the calculation regions that are not focally amplified (CN < 4.5 default), as well as redundant filter regions that are also present in the low-complexity or low-mappability database used by AmpliconArchitect.

In addition to the metrics output by the amplicon similarity script, we also calculated the amplicon length coverage percentage. This metric was computed as 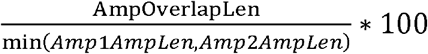 to determine how well the amplicons overlap between a pair covering the amplicon with the lower length.

## Statistical methods

Statistical tests, test statistics, and p-values are indicated where appropriate in the main text. Categorical associations were established using the chi-square test of independence if N>5 for all categories; otherwise, the Fisher’s exact test was used. For both tests, the Python package scipy.stats v1.5.3 implementation was used [^27^]. In addition, statistical analysis for the paired tumors was performed using the McNemar test [^28^] implemented in statsmodels v0.12.0 [^29^].

### Single cell data processing and clustering

Sequencing data for both tumor-PDX pairs (RCMB56-ht/pdx and RC124-ht/pdx) were uniformly processed using CellRanger ARC (v2.0.0 for RCMB56, v2.0.2 for RC124) with default parameters, followed by Seurat (v4.0.4 for RCMB56, v5.3.0 for RC124). For RCMB56, cell barcodes were retained based on quality thresholds: ATAC mitochondrial fraction <0.1, ATAC read count between 1,000-70,000, and RNA read count between 500 (ht) or 1000 (pdx) and 25,000. Doublets were identified and removed using DoubletFinder v2.0 (RRID:SCR_018771) with default parameters. Host cells were removed from PDX samples using Xenocell v1.0.1, filtering barcodes with < 90% uniquely mapped reads to the human genome. Following preprocessing, 2,986 RCMB56-ht and 10,400 RCMB56-pdx cells remained, whereas 5,669 RC124-ht and 6,338 RC124-pdx cells were present after applying similar filtering criteria.

Clustering was performed independently for each sample using the weighted nearest neighbor algorithm [^30^] with default parameters. Cell-type identities were assigned by identifying differentially expressed genes (DEGs) for each cluster using Seurat’s FindAllMarkers function and cross-referencing against known cell type marker genes [^31^]. Only primary tumor clusters (RCMB56-ht and RC124-ht) were labeled in this manner. For RC124 copy number analysis, ecDNA-quant outputs (see below) were additionally utilized for cell type identification.

### Identification of ecDNA-containing cells

ecDNA-containing cells were identified by permutation tests comparing scATAC-seq read coverage at the ecDNA locus, using AmpliconArchitect on bulk WGS, as described above, to read coverage elsewhere in the genome. This code is available at https://github.com/auberginekenobi/ecdna-quant. Briefly, deduplicated scATAC-seq reads were obtained from the fragments.tsv output of CellRanger ARC and sorted by barcode. For Monte Carlo permutation testing, 1000 random contiguous regions of the genome, excluding centromeres, telomeres, known ecDNAs, and low-mappability regions, were generated using bedtools v2.27.1[^32^]. Read coverage was counted using PyRanges v0.0.112 [^33^] and scaled to region length. For each cell, empirical p-values were estimated as 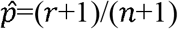 [^34^]. Multiple hypothesis correction was performed using the Benjamini-Hochberg correction as described above. Z-scores were calculated using the standard formula, comparing the average read coverage at the ecDNA-amplified region to the mean and variance of the Monte Carlo permutations.

### Single-cell genomic copy number estimation

To investigate copy number aberrations based on single-nucleus ATAC-sequencing data, we used an adapted version of InferCNV as described in Okonechnikov et al [^35^].

## Author’s Contributions

**R. Kenkre:** Conceptualization, data aggregation, formal analysis, validation, investigation, writing-original draft, writing-review, and editing. **Jon Larson:** Conceptualization, methodology, writing-review, and editing. **O. Chapman:** Conceptualization, data aggregation, formal analysis, writing-original draft, writing-review, and editing. **J. Luebeck:** Conceptualization, methodology, writing-review, and editing. **Y.Y. Lo:** Conceptualization, methodology, writing-review and editing. **M. Paul:** Validation, writing–review, and editing. **W. Zhang:** Data aggregation, formal analysis. **V. Bafna:** Supervision, validation, writing–review, and editing. **R.J. Wechsler-Reya:** Supervision, validation, writing–review, and editing. **L. Chavez:** Conceptualization, resources, supervision, funding acquisition, writing–review, and editing.

## Acknowledgements

This work was supported by a generous endowment by the Clayes Foundation to the Research Center for Neuro-Oncology and Genomics within the Rady Children’s Institute for Genomic Medicine, a Hannah’s Heroes St. Baldrick’s Scholar Award (L.C.), the Dragon Master Foundation (L.C.), funding from the National Institutes of Health (NIH) National Institute of Neurological Disorders and Stroke Institute R01 NS132780 (L.C.), the NIH National Cancer Institute P30 CA030199 (L.C.), and F31 CA271777 (O.S.C.). We would like to thank the Moores Cancer Center Pilot Grant (L.C., V.B., and J.P.M.). This work used Expanse cluster computing services at the San Diego Supercomputer Center through allocation BIO210026 from the Advanced Cyberinfrastructure Coordination Ecosystem: Services & Support (ACCESS) program, which is supported by National Science Foundation grants #2138259, #2138286, #2138307, #2137603, and #2138296. This research was conducted using data made available by The Children’s Brain Tumor Network (formerly the Children’s Brain Tumor Tissue Consortium) and the St. Jude Cloud. We thank J. H. Zhang, C. McLeod, D. Miller, B. Curran, and A. Resnick for facilitating data access. Support through grant P30 CA030199 to the Animal Resources core facility at Sanford Burnham Prebys (NCI designated Cancer Center) is gratefully acknowledged.

## Figure Legends

**Supplementary Figure 1: Visualizations of ecDNA genomic regions amongst human tumor-PDX pairs that transitioned from ecDNA-positive to ecDNA-negative status**.

Integrative Genomics Viewer (IGV) visualizations of the three tumor-PDX pairs for which ecDNA was lost. Regions with high coverage indicate potential ecDNA locations. The red horizontal line indicates predicted ecDNA-amplified location. _X indicates PDX model and any other annotation indicates human tumor. ecDNA-amplified gene FKBP5 in patient **(a)** SJRB046160; no ecDNA-amplified genes in the other two primary tumors for patients **(b)** SJRHB013758 and **(c)** SJRHB063825.

**Supplementary Figure 2: Visualizations of ecDNA genomic regions amongst human tumor-PDX pairs that transitioned from ecDNA-negative to ecDNA-positive status**.

Integrative Genomics Viewer (IGV) visualizations of the three tumor-PDX pairs for which ecDNA was lost. Regions of high coverage indicate potential ecDNA locations. Red horizontal line indicates predicted ecDNA-amplified location. _X indicates PDX model and any other annotation indicates human tumor. **(a)** Patient SJNBL105 (neuroblastoma), **(b)** Patient SJNBL012407 (neuroblastoma), **(c)** Patient SJNBL046145 (neuroblastoma), **(d)** Patient SJOS012409 (osteosarcoma).

**Supplementary Figure 3: Amplicon similarity score metric distribution**.

**(a)** Boxplot showing amplicon similarity score distribution of ecDNA-positive tumor-PDX pairs.

**(b)** Boxplot showing genomic segment score distribution of ecDNA-positive tumor-PDX pairs.

**(c)** Boxplot showing breakpoint sharing score distribution of ecDNA-positive tumor-PDX pairs.

**Supplementary Figure 4: Amplicon graphs of ecDNA-positive human tumor-PDX pairs**.

**(a)** Osteosarcoma pair (SJOS016016) demonstrating perfect sequence conservation with similarity score 1.0. **(b)** Osteosarcoma pair (SJOS001121) with amplicon similarity score 0.84. **(c) N**euroblastoma pair (SJNBL124) with amplicon similarity score 0.74. **(d)** Neuroblastoma pair (SJNBL046148) demonstrating with amplicon similarity score 0.37.

**Supplementary Figure 5: Joint single-cell clustering of RCMB56 tumor-PDX pair**

Conos clustering of the transcriptome profile in RCMB56-PDX, overlaid on the transcriptome profile of RCMB56-HT to highlight similarities and differences in gene expression patterns between the two samples.

**Supplementary Table 1: Sample Information of St Jude PDX models and Rady PDX models**.

**(a)** Includes St Jude PDX model information such as sample name, sample type, cancer type, sex, ethnicity, and race. **(b)** Includes Rady PDX model information such as sample name, sample type, cancer type, sex, ethnicity, and race.

**Supplementary Table 2: Amplicon classifier results of St Jude PDX models and paired human tumors**.

**(a)** Shows information such as amplicon classification and oncogenes for each PDX sample.

**(b)** Shows information such as amplicon classification and oncogenes for each paired human tumor.

**(c)** Shows information such as amplicon classification and oncogenes for each Rady MB tumor-PDX pairs.

**Supplementary Table 3: Paired ecDNA Status Results of PDX models and Human Tumors of Origin**.

Paired tumor table displaying ecDNA status and corresponding cancer type.

**Supplementary Table 4: Paired Tumors with ecDNA Gains in PDX Models and Corresponding Genes**

Paired tumor table showing only ecDNA being gained in PDX models, corresponding tumor type, and ecDNA-amplified genes.

**Supplementary Table 5: Amplicon Similarity Results of ecDNA+ human tumor-PDX pairs from Amplicon Suite**

**(a)** Amplicon similarity results of St Jude ecDNA-positive human tumor-PDX pairs. **(b)**

Amplicon similarity results of Rady ecDNA-positive human tumor-PDX pairs.

**Supplementary Table 6:** Copy Number Comparison of ecDNA-amplified Genes within Paired Tumors

