## Supplementary figures and images for "Preservation and Clonal Behavior of Extrachromosomal DNA in Patient-Derived Xenograft Models of Childhood Cancers"

### Supplementary Figure 1

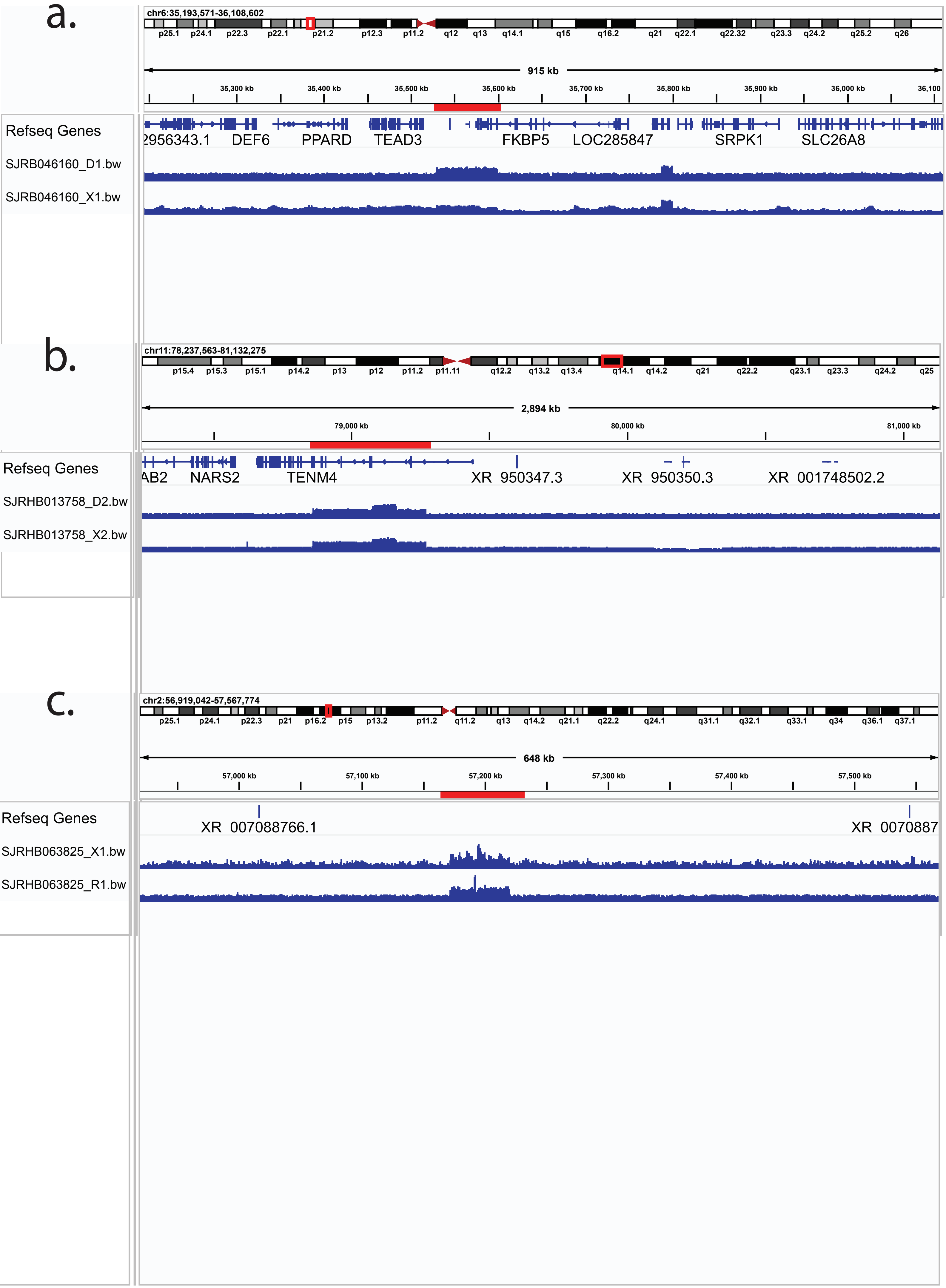


Supplementary Figure 1

### Supplementary Figure 2

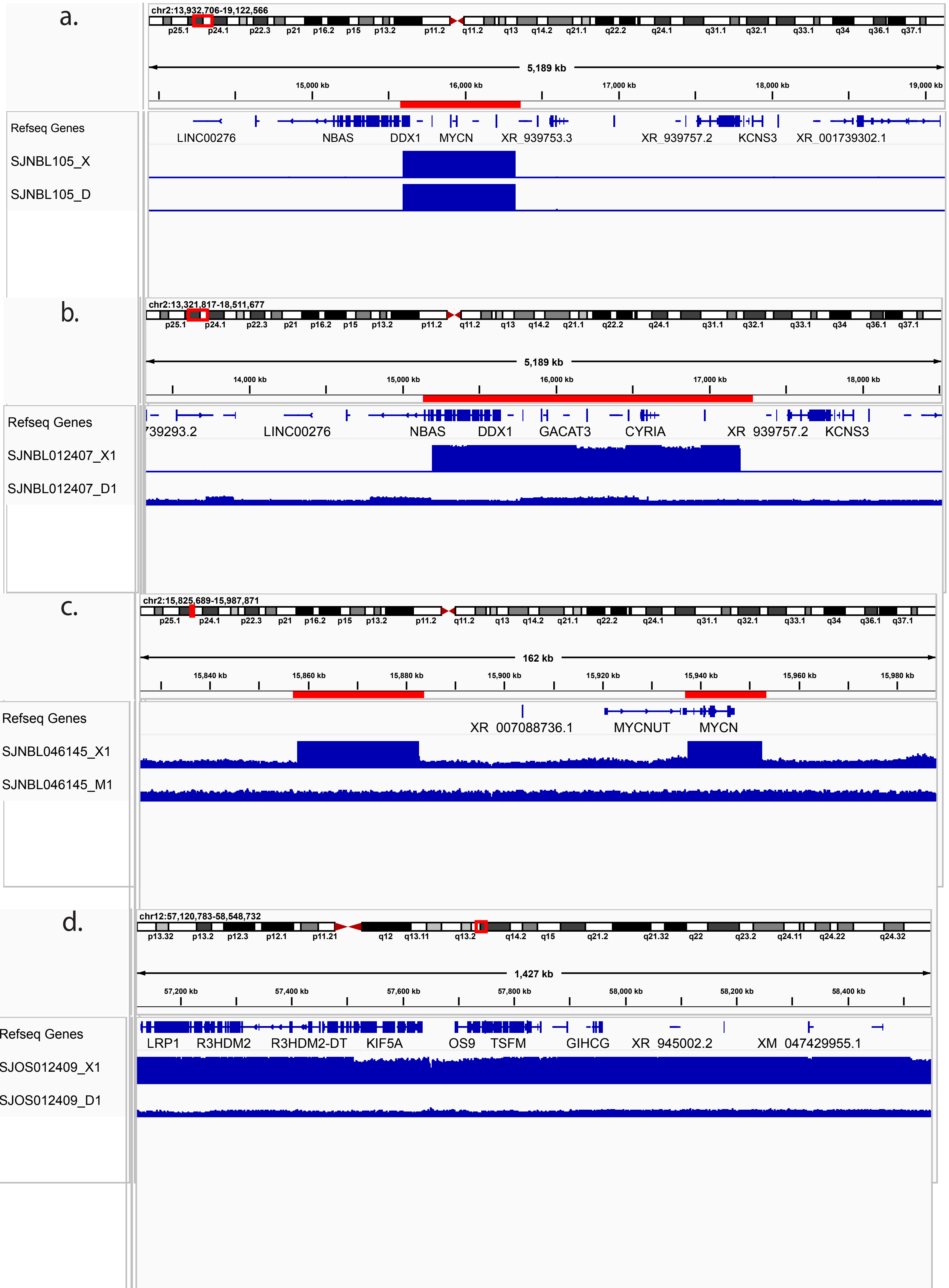


Supplementary Figure 2

### Supplementary Figure 3

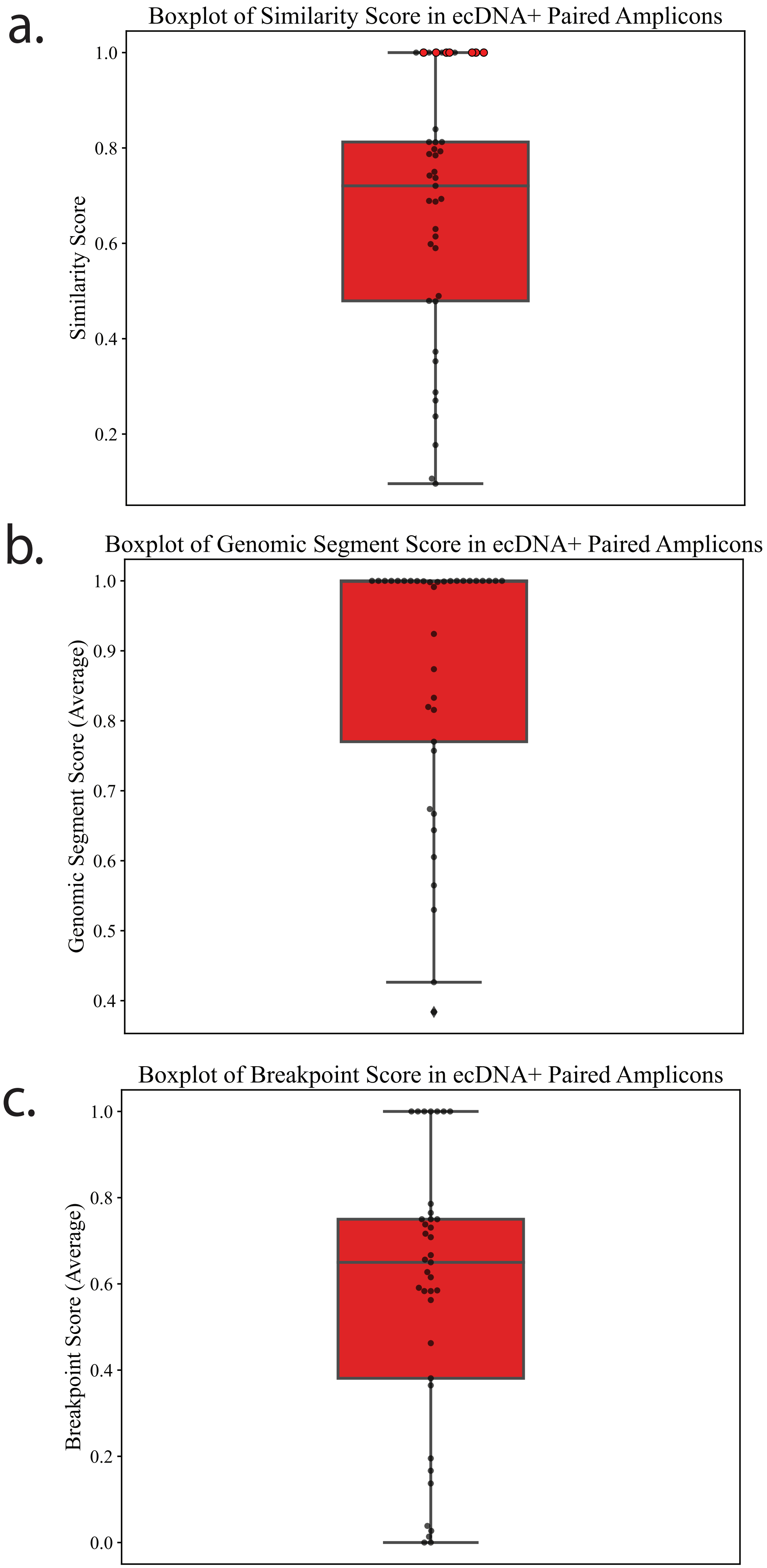


Supplementary Figure 3

### Supplementary Figure 4

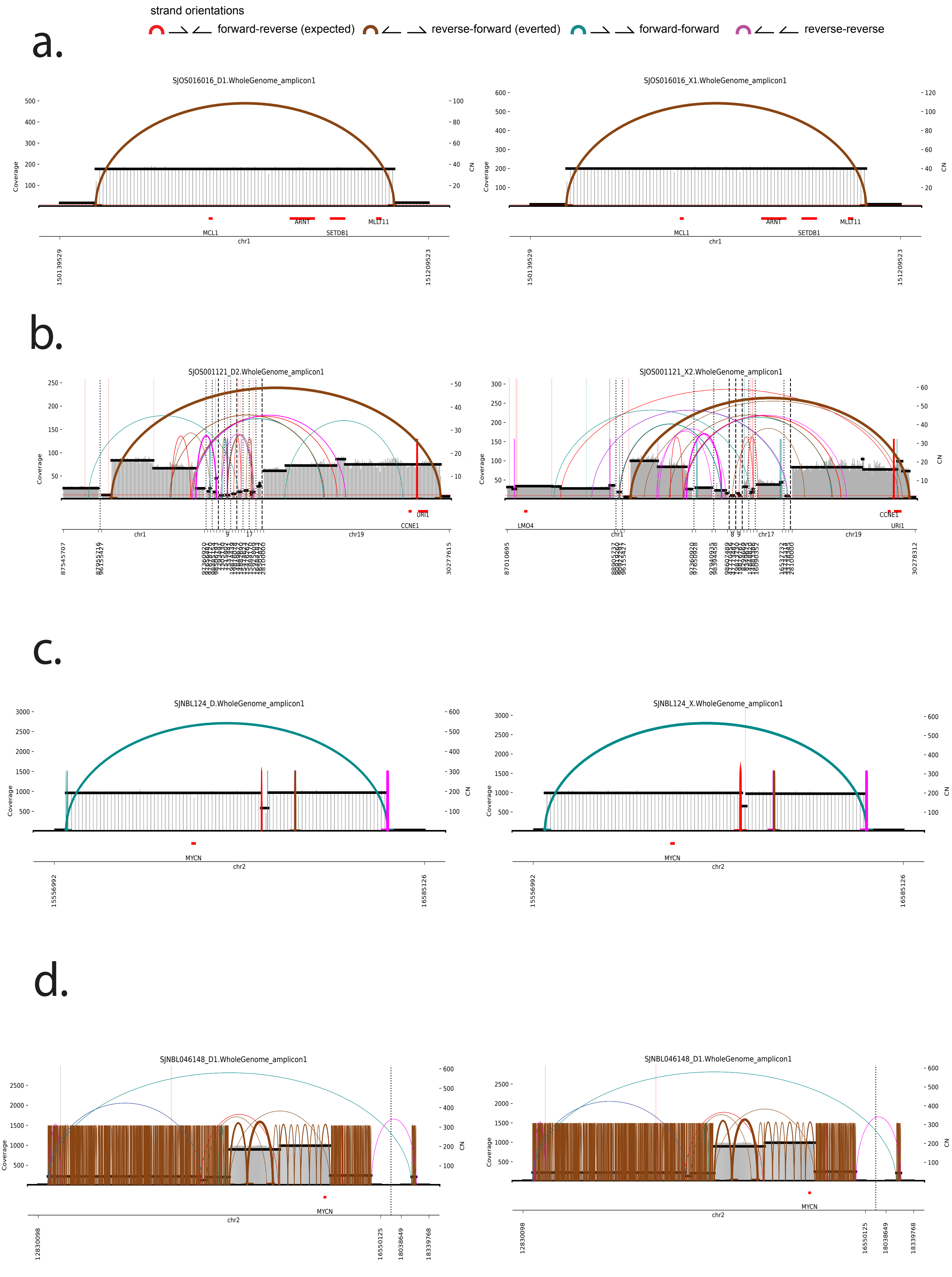


Supplementary Figure 4

### Supplementary Figure 5

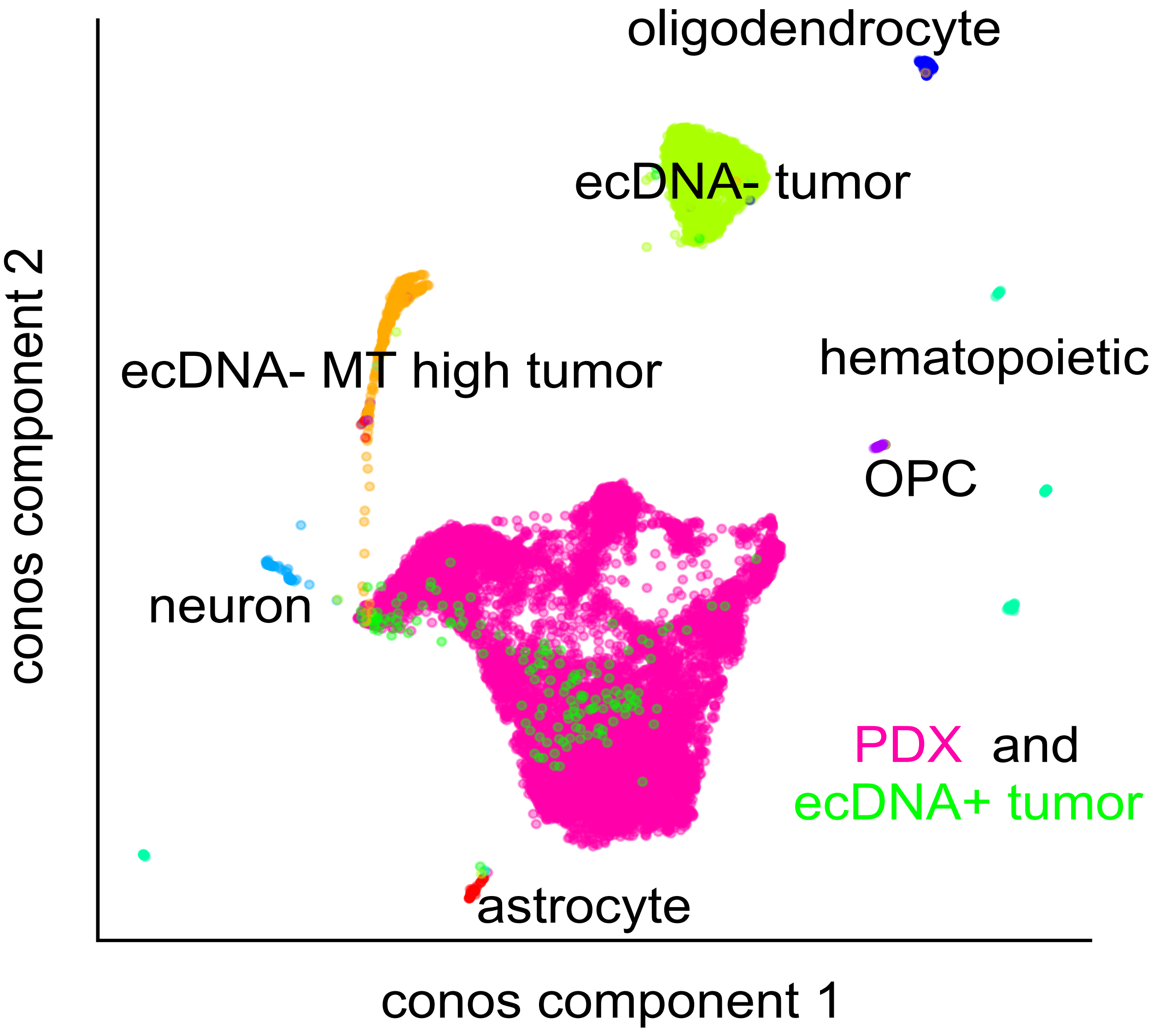


Supplementary Figure 5
